# Is it actionable? An Evaluation of the Rapid PCR-Based Blood Culture Identification Panel on the Management of Gram-Positive and Gram-Negative Blood Stream Infections

**DOI:** 10.1101/375477

**Authors:** Andrew S. Tseng, Sabirah N. Kasule, Felicia Rice, Lanyu Mi, Lynn Chan, Maria T. Seville, Thomas E. Grys

## Abstract

**Background:** There is growing interest in the use of rapid blood culture identification (BCID) panels in antimicrobial stewardship programs (ASP). While many studies have looked at its clinical and economic utility, its comparative utility in gram-positive and gram-negative blood stream infections (BSI) have not been as well characterized.

**Methods:** The study was a quasi-experimental retrospective study at the Mayo Clinic in Phoenix, Arizona. All adult patients with positive blood cultures before BCID implementation (June 2015 to December 2015) and after BCID implementation (June 2016 to December 2016) were included. The outcomes of interest included: time to first appropriate antibiotic escalation, time to first appropriate antibiotic de-escalation, time to organism identification, LOS, infectious disease consultation, discharge disposition, and in-hospital mortality.

**Results:** In total, 203 patients were included in this study. There was a significant difference in the time to organism identification between pre- and post-BCID cohorts (27.1h vs. 3.3h, p<0.0001). BCID did not significantly reduce the time to first appropriate antimicrobial escalation or de-escalation for either GP-BSIs or GN-BSIs. Providers were more likely to escalate antimicrobial therapy in GP-BSIs after gram stain and more likely to de-escalate therapy in GN-BSIs after susceptibilities. While there were no significant differences in changes in antimicrobial therapy after organism identification by BCID, over a quarter of providers (28.1%) made changes after organism identification.

**Conclusions:** While BCID significantly reduced the time to identification for both GP-BSIs and GN-BSIs, BCID did not reduce the time to first appropriate antimicrobial escalation and de-escalation.

## INTRODUCTION

Blood stream infections (BSI) are life-threatening events which require effective treatment for optimal outcomes. Often, patients are on multiple antimicrobial drugs until the offending pathogen is identified. Thus, shortening the time to identification (ID) and antimicrobial susceptibility testing (AST) is essential to reduce exposure to unnecessary antimicrobial drugs (1). Rapid blood culture identification (BCID) panels provide an opportunity to improve use of antimicrobial drugs and improve patient outcomes. These BCID panels are also a new tool for effective implementation of antimicrobial stewardship programs (ASP) with active surveillance and proactive intervention by an infectious disease specialist or pharmacist.

Observational studies have shown that rapid organism identification in BSI was associated with a decrease in mortality, length of stay (LOS) and cost (2-10). These benefits are largely derived from appropriate antimicrobial escalation, timely antimicrobial de-escalation and utilization of narrow-spectrum antimicrobials resulting in shorter lengths of stays, less treatment of contaminant blood cultures and reduced cost of antibiotics. A large single-center prospective study confirmed the utility of the rapid PCR-based BCID panels in reducing unnecessary antibiotic use, particularly in conjunction with ASP. However, the investigators noted that the median time to appropriate de-escalation in the BCID-only group was not significantly different than control (no intervention), and only with the addition of proactive ASP to the BCID was there a significant decrease in median time to appropriate de-escalation (11). This was attributed to the likely scenario that, even with rapid organism identification, providers were hesitant to de-escalate antimicrobial therapies without antimicrobial susceptibility testing (AST) results unless they received specific guidance from ASP infectious disease specialists or pharmacists. Another possibility is that in the absence of proactive reporting of the identification, the provider may not review or act upon the new result until the following day.

Currently, many BCID panels utilize gene testing to determine antimicrobial resistance in certain gram-positive organisms, such as the mecA, vanA and vanB genes for methicillin and vancomycin resistance in staphylococci and enterococci, and in certain gram-negative organisms, including the bla_KPC_ gene for carbapenem resistance in *Enterobacteriaceae* (particularly *Escherichia coli* and *Klebsiella pneumoniae)* (12). While the presence of these genes may allow for appropriate escalation of antimicrobial therapy, clinicians are often hesitant to de-escalate before susceptibility testing, since there are myriad genotypic mutations that may confer phenotypic resistance, which were not included in the BCID panel. The purpose of laboratory testing is to provide actionable information in a rapid manner to improve patient care. The purpose of the present study is to evaluate the clinical utility of BCID panels in gram-positive versus gram-negative BSI.

## MATERIALS AND METHODS

### Study Design

The study was a quasi-experimental retrospective study at a medium-sized (268 inpatient bed) academic tertiary referral hospital with a high volume of solid and bone marrow transplant patients. All adult (>18 years of age) patients with positive blood cultures before BCID implementation (June 2015 to December 2015) and after BCID implementation (June 2016 to December 2016) were included. Exclusion criteria included non-bacterial BSI, contaminants, mixed gram-positive/gram-negative BSI, expiration or hospice enrollment prior to hospitalization, history of previously positive blood culture with the same organism within 90 days and any patients who were not admitted to our institution or had positive blood cultures obtained as an outpatient. When the laboratory deployed the BCID test, there was a concurrent implementation of a modest, proactive ASP. The laboratory would email members of the ASP group, who would verify that the patient was on effective therapy, and make recommendations to the providers if there were opportunities to de-escalate therapy. The ASP team would review the positive BCID results during normal working hours, and generally same-day, or the next business day if the BCID result was sent after hours or on weekends. Escalation in antimicrobial therapy was defined as the addition of an additional antibiotic or substitution of an antibiotic with broader coverage. De-escalation was defined as the removal of an antibiotic or substitution of an antibiotic with narrower coverage. Changes that were neither escalation nor de-escalation were substitutions of antibiotics to cover completely different organisms (e.g. switching from vancomycin to piperacillin-tazobactam after gram stain showed gram-negative bacillus). Notably, the ASP approach was implemented for routine care and not specifically for this study. The study was approved by the Mayo Clinic Institutional Review Board.

### BCID Panel

Our institution utilized the FilmArray BCID (BioFire Diagnostics, LLC, Salt Lake City, UT), which was performed on all positive blood cultures beginning on June 1^st^, 2016. The BCID can identify *Staphylococcus spp., S. aureus, Streptococcus spp., S. agalactiae, S. pyogenes, S. pneumoniae, Enterococcus spp., Listeria monocytogenes, Enterobacteriaceae, Escherichia coli, Enterobactercloacae complex, Klebsiellaoxytoca, K. pneumoniae, Serratia spp., Proteus spp., Acinetobacter baumannii, Haemophilus influenzae, Neisseria meningitidis, Pseudomonas aeruginosa, Candida albicans, C. glabrata, C. krusei, C. parapsilosis,* and *C. tropicalis* and 4 antibiotic resistance genes, mecA, vanA and vanB (vanA/B), and bla_K_pc.

### Outcome

The outcomes of interest included: (1) time to first appropriate antibiotic escalation (initiation of broader-spectrum antibiotics), (2) time to first appropriate antibiotic de-escalation (alteration to narrow-spectrum antibiotics), (3) time to organism identification (the time of organism identification either by conventional testing or BCID from the time of culture positivity), (4) LOS, (5) infectious disease consultation and (6) patient disposition (home, facility, hospice or death) at discharge. Other metrics included the antimicrobial therapy adjustments following each stage of BSI investigation (GS, ID, AST).

### Statistical Analysis

Continuous data were presented as medians with interquartile ranges, unless otherwise specified. Categorical data were presented as frequencies and percentages. Data were assessed using graphical and descriptive analysis to evaluate the distributions and assess for outliers. Wilcoxon rank sum test was used for continuous variables. Fisher’s exact test or Chi-squared test was used for categorical variables. Results were considered statistically significant at a (2-tailed) P-value of < 0.05. All statistical analyses were completed using SAS Studio 3.7 (SAS Institute Inc, Cary, North Carolina, USA).

## RESULTS

### Patient Characteristics and Co-morbidities

There were 381 positive blood cultures identified in the two aforementioned time periods pre- and post-BCID (118 and 137 positive blood cultures, respectively). After exclusion of duplicates, contaminants and non-inpatient encounters, 203 patients (mean age 65.0±15.2, 35.0% female) were included. The study CONSORT diagram showing the derivation of the study cohort is shown in Figure 1. For most demographic and medical co-morbidity variables, there were no significant differences between gram-negative and gram-positive BSIs. The data is summarized in Table 1.

**Figure 1:**
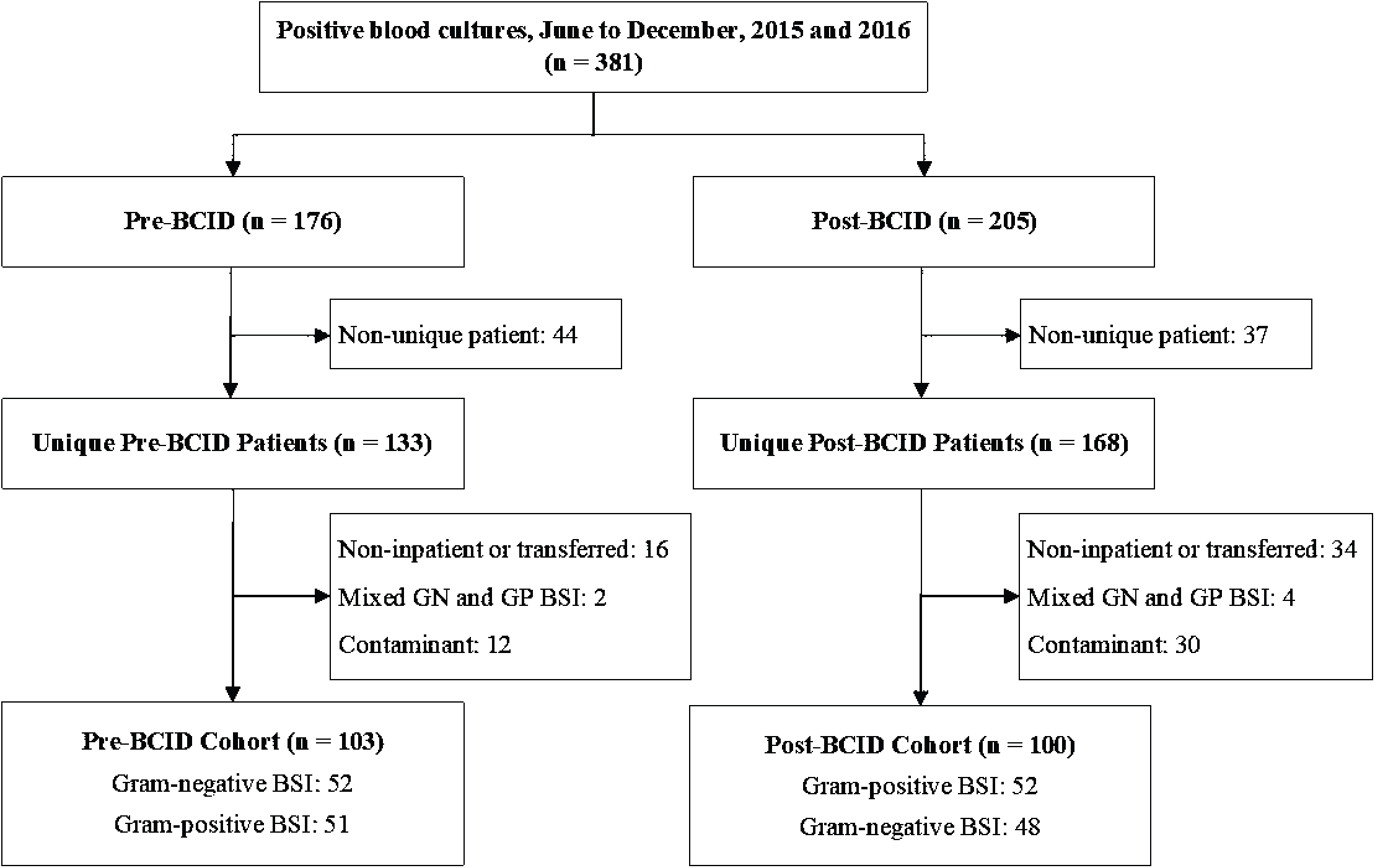
Study CONSORT diagram. CONSORT diagram for the derivation of the final cohort.

**Figure 2:**
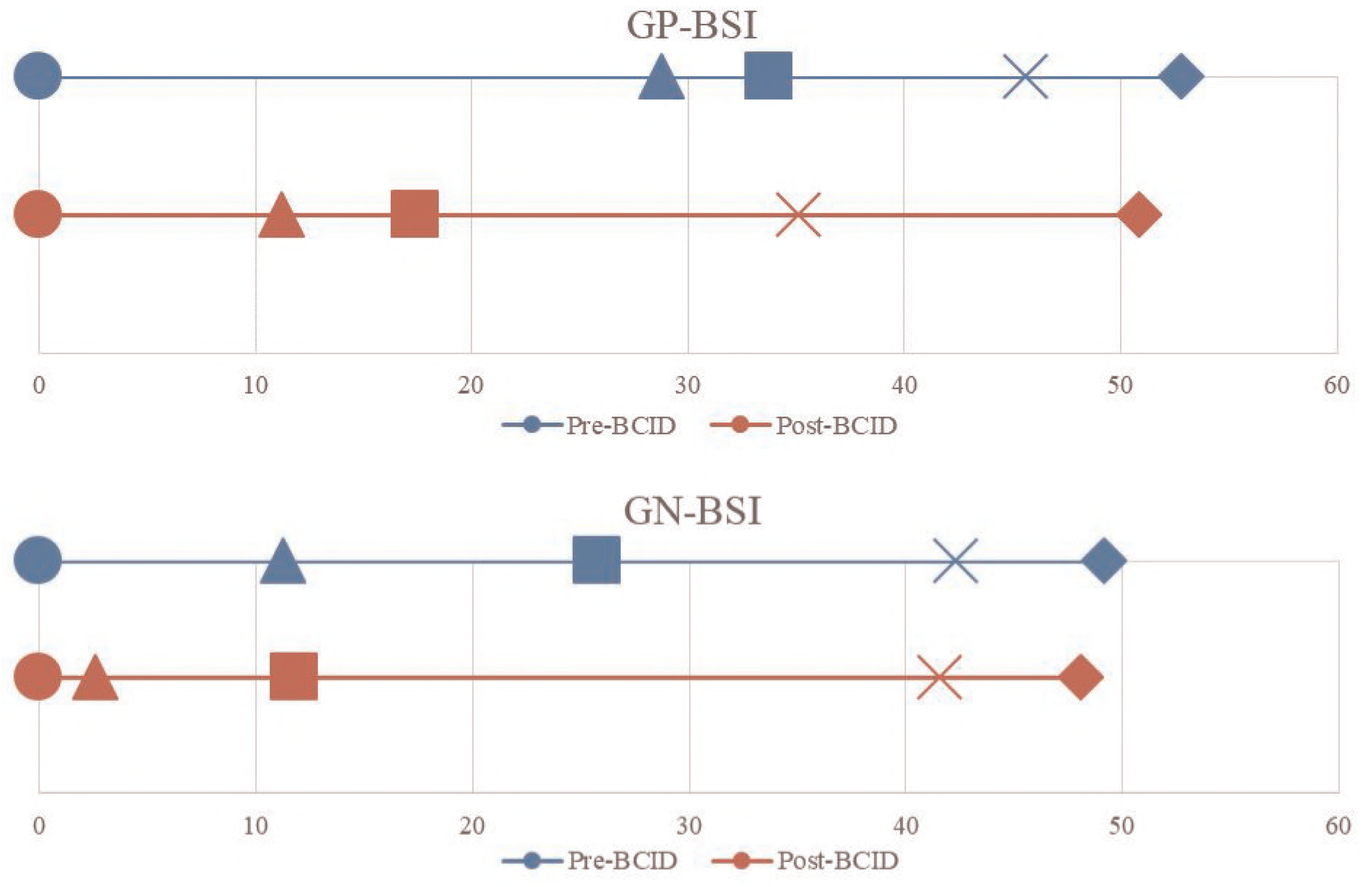
Timeline of key microbiological and clinical events from time to positivity and gram stain. The timeline of key events include time to positive and gram stain (circle), time to first escalation (triangle), time to identification (square), time to first de-escalation (x), and time to susceptibilities (diamond). Times are represented in hours. There were no significant differences in any of the time points for each key event for pre-versus post-BCID by gram stain.?

**Table 1:**
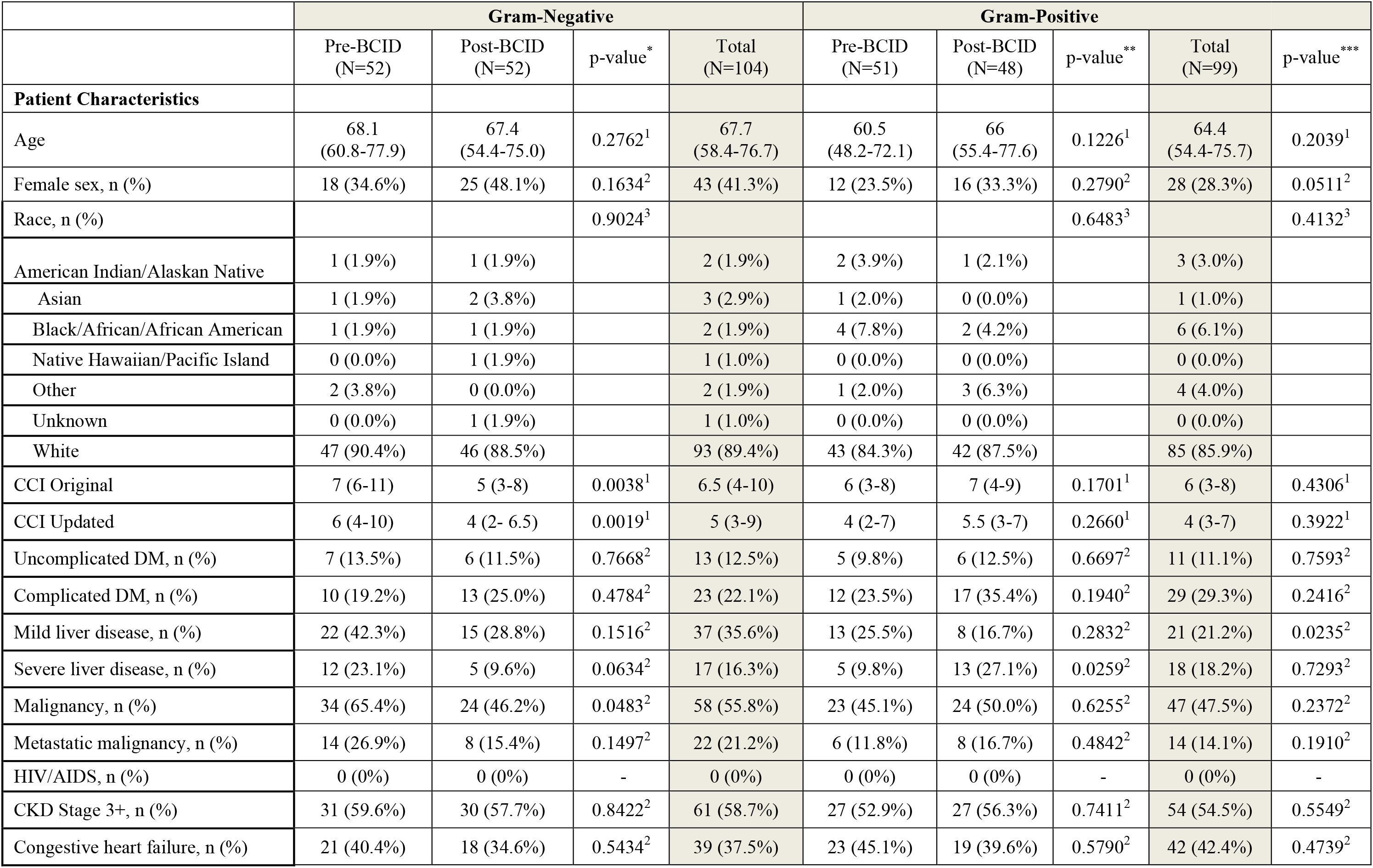
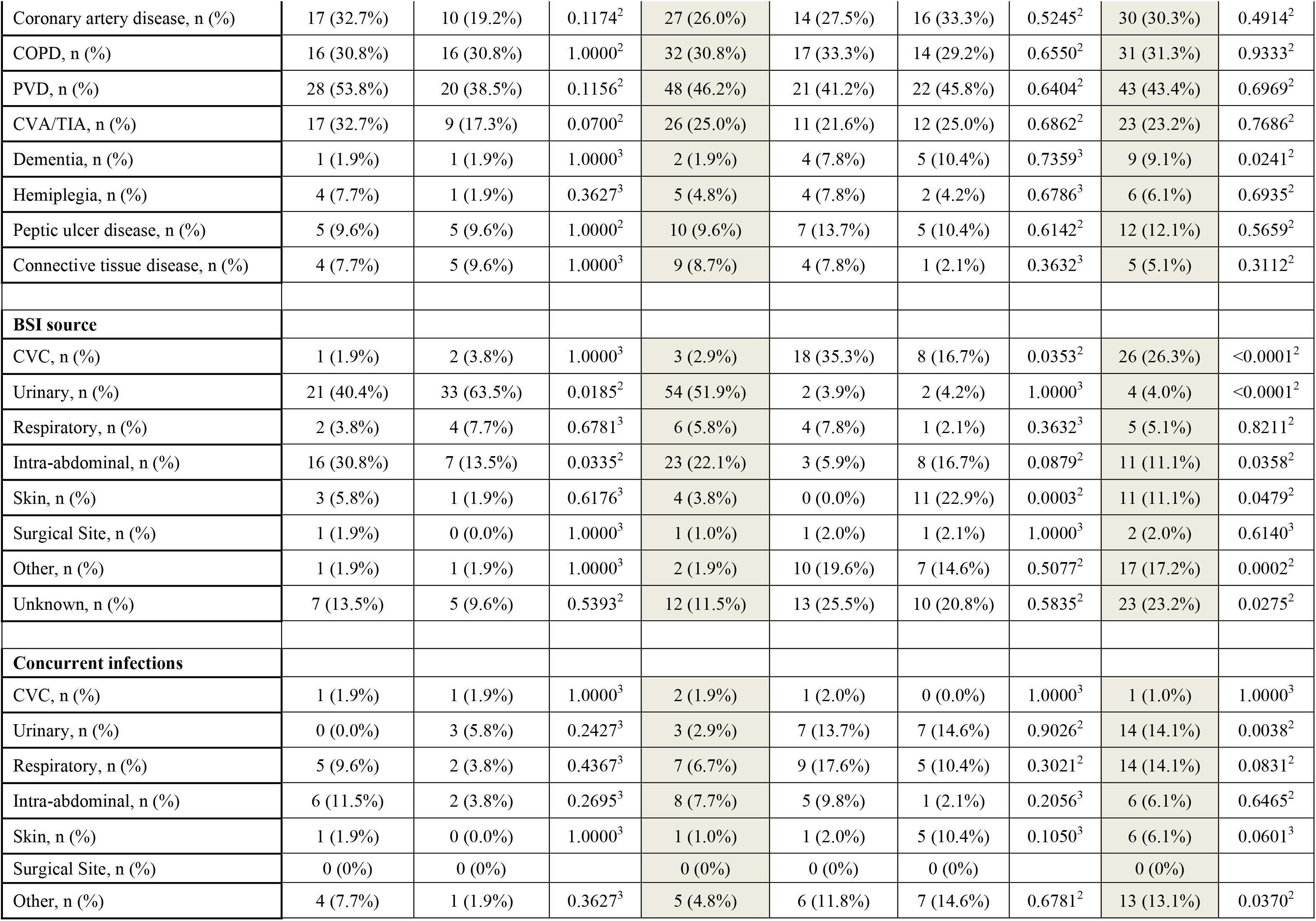
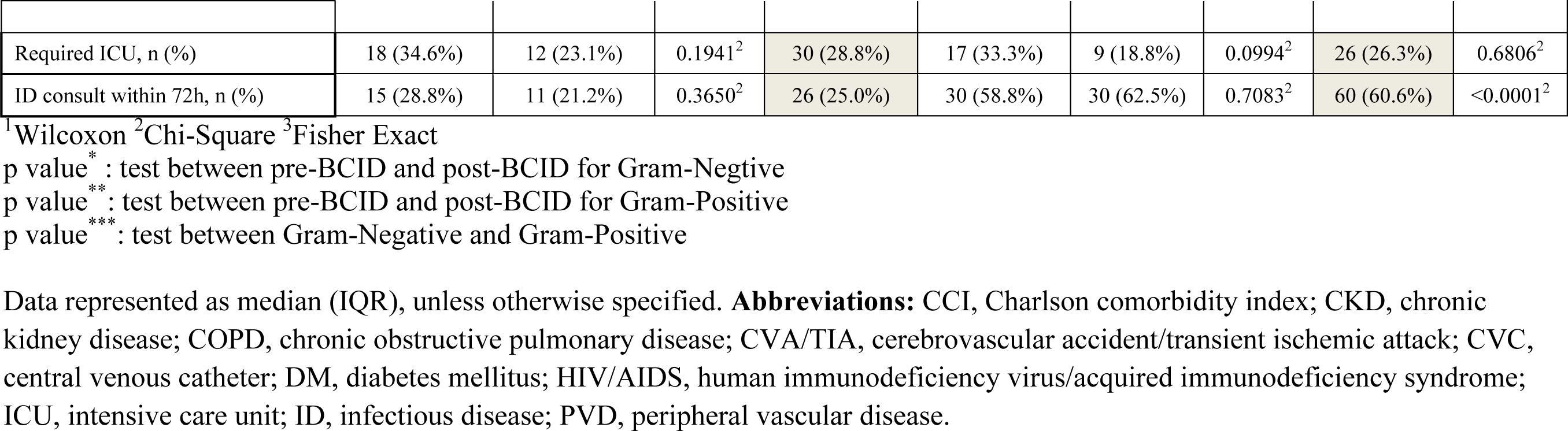
Patient Baseline and BSI Characteristics

|  | Gram-Negative |  |  |  | Gram-Positive |  |  |  |  |
| --- | --- | --- | --- | --- | --- | --- | --- | --- | --- |
|  | Pre-BCID<br>(N=52) | Post-BCID<br>(N=52) | p-value * | Total<br>(N=104) | Pre-BCID<br>(N=51) | Post-BCID<br>(N=48) | p-value ** | Total<br>(N=99) | p-value *** |
| **Patient Characteristics** |  |  |  |  |  |  |  |  |  |
| Age | 68.1<br>(60.8-77.9) | 67.4<br>(54.4-75.0) | 0.2762<sup>1</sup> | 67.7<br>(58.4-76.7) | 60.5<br>(48.2-72.1) | 66<br>(55.4-77.6) | 0.1226<sup>1</sup> | 64.4<br>(54.4-75.7) | 0.2039<sup>1</sup> |
| Female sex, n (%) | 18 (34.6%) | 25 (48.1%) | 0.1634<sup>2</sup> | 43 (41.3%) | 12 (23.5%) | 16 (33.3%) | 0.2790<sup>2</sup> | 28 (28.3%) | 0.0511<sup>2</sup> |
| Race, n (%) |  |  | 0.9024<sup>3</sup> |  |  |  | 0.6483<sup>3</sup> |  | 0.4132<sup>3</sup> |
| American Indian/Alaskan Native | 1 (1.9%) | 1 (1.9%) |  | 2 (1.9%) | 2 (3.9%) | 1 (2.1%) |  | 3 (3.0%) |  |
| Asian | 1 (1.9%) | 2 (3.8%) |  | 3 (2.9%) | 1 (2.0%) | 0 (0.0%) |  | 1 (1.0%) |  |
| Black/African/African American | 1 (1.9%) | 1 (1.9%) |  | 2 (1.9%) | 4 (7.8%) | 2 (4.2%) |  | 6 (6.1%) |  |
| Native Hawaiian/Pacific Island | 0 (0.0%) | 1 (1.9%) |  | 1 (1.0%) | 0 (0.0%) | 0 (0.0%) |  | 0 (0.0%) |  |
| Other | 2 (3.8%) | 0 (0.0%) |  | 2 (1.9%) | 1 (2.0%) | 3 (6.3%) |  | 4 (4.0%) |  |
| Unknown | 0 (0.0%) | 1 (1.9%) |  | 1 (1.0%) | 0 (0.0%) | 0 (0.0%) |  | 0 (0.0%) |  |
| White | 47 (90.4%) | 46 (88.5%) |  | 93 (89.4%) | 43 (84.3%) | 42 (87.5%) |  | 85 (85.9%) |  |
| CCI Original | 7 (6-11) | 5 (3-8) | 0.0038<sup>1</sup> | 6.5 (4-10) | 6 (3-8) | 7 (4-9) | 0.1701<sup>1</sup> | 6 (3-8) | 0.4306<sup>1</sup> |
| CCI Updated | 6 (4-10) | 4 (2- 6.5) | 0.0019<sup>1</sup> | 5 (3-9) | 4 (2-7) | 5.5 (3-7) | 0.2660<sup>1</sup> | 4 (3-7) | 0.3922<sup>1</sup> |
| Uncomplicated DM, n (%) | 7 (13.5%) | 6 (11.5%) | 0.7668<sup>2</sup> | 13 (12.5%) | 5 (9.8%) | 6 (12.5%) | 0.6697<sup>2</sup> | 11 (11.1%) | 0.7593<sup>2</sup> |
| Complicated DM, n (%) | 10 (19.2%) | 13 (25.0%) | 0.4784<sup>2</sup> | 23 (22.1%) | 12 (23.5%) | 17 (35.4%) | 0.1940<sup>2</sup> | 29 (29.3%) | 0.2416<sup>2</sup> |
| Mild liver disease, n (%) | 22 (42.3%) | 15 (28.8%) | 0.1516<sup>2</sup> | 37 (35.6%) | 13 (25.5%) | 8 (16.7%) | 0.2832<sup>2</sup> | 21 (21.2%) | 0.0235<sup>2</sup> |
| Severe liver disease, n (%) | 12 (23.1%) | 5 (9.6%) | 0.0634<sup>2</sup> | 17 (16.3%) | 5 (9.8%) | 13 (27.1%) | 0.0259<sup>2</sup> | 18 (18.2%) | 0.7293<sup>2</sup> |
| Malignancy, n (%) | 34 (65.4%) | 24 (46.2%) | 0.0483<sup>2</sup> | 58 (55.8%) | 23 (45.1%) | 24 (50.0%) | 0.6255<sup>2</sup> | 47 (47.5%) | 0.2372<sup>2</sup> |
| Metastatic malignancy, n (%) | 14 (26.9%) | 8 (15.4%) | 0.1497<sup>2</sup> | 22 (21.2%) | 6 (11.8%) | 8 (16.7%) | 0.4842<sup>2</sup> | 14 (14.1%) | 0.1910<sup>2</sup> |
| HIV/AIDS, n (%) | 0 (0%) | 0 (0%) | - | 0 (0%) | 0 (0%) | 0 (0%) | - | 0 (0%) | - |
| CKD Stage 3+, n (%) | 31 (59.6%) | 30 (57.7%) | 0.8422<sup>2</sup> | 61 (58.7%) | 27 (52.9%) | 27 (56.3%) | 0.7411<sup>2</sup> | 54 (54.5%) | 0.5549<sup>2</sup> |
| Congestive heart failure, n (%) | 21 (40.4%) | 18 (34.6%) | 0.5434<sup>2</sup> | 39 (37.5%) | 23 (45.1%) | 19 (39.6%) | 0.5790<sup>2</sup> | 42 (42.4%) | 0.4739<sup>2</sup> |
| Coronary artery disease, n (%) | 17 (32.7%) | 10 (19.2%) | 0.1174<sup>2</sup> | 27 (26.0%) | 14 (27.5%) | 16 (33.3%) | 0.5245<sup>2</sup> | 30 (30.3%) | 0.4914<sup>2</sup> |
| COPD, n (%) | 16 (30.8%) | 16 (30.8%) | 1.0000<sup>2</sup> | 32 (30.8%) | 17 (33.3%) | 14 (29.2%) | 0.6550<sup>2</sup> | 31 (31.3%) | 0.9333<sup>2</sup> |
| PVD, n (%) | 28 (53.8%) | 20 (38.5%) | 0.1156<sup>2</sup> | 48 (46.2%) | 21 (41.2%) | 22 (45.8%) | 0.6404<sup>2</sup> | 43 (43.4%) | 0.6969<sup>2</sup> |
| CVA/TIA, n (%) | 17 (32.7%) | 9 (17.3%) | 0.0700<sup>2</sup> | 26 (25.0%) | 11 (21.6%) | 12 (25.0%) | 0.6862<sup>2</sup> | 23 (23.2%) | 0.7686<sup>2</sup> |
| Dementia, n (%) | 1 (1.9%) | 1 (1.9%) | 1.0000<sup>3</sup> | 2 (1.9%) | 4 (7.8%) | 5 (10.4%) | 0.7359<sup>3</sup> | 9 (9.1%) | 0.0241<sup>2</sup> |
| Hemiplegia, n (%) | 4 (7.7%) | 1 (1.9%) | 0.3627<sup>3</sup> | 5 (4.8%) | 4 (7.8%) | 2 (4.2%) | 0.6786<sup>3</sup> | 6 (6.1%) | 0.6935<sup>2</sup> |
| Peptic ulcer disease, n (%) | 5 (9.6%) | 5 (9.6%) | 1.0000<sup>2</sup> | 10 (9.6%) | 7 (13.7%) | 5 (10.4%) | 0.6142<sup>2</sup> | 12 (12.1%) | 0.5659<sup>2</sup> |
| Connective tissue disease, n (%) | 4 (7.7%) | 5 (9.6%) | 1.0000<sup>3</sup> | 9 (8.7%) | 4 (7.8%) | 1 (2.1%) | 0.3632<sup>3</sup> | 5 (5.1%) | 0.3112<sup>2</sup> |
| **BSI source** |  |  |  |  |  |  |  |  |  |
| CVC, n (%) | 1 (1.9%) | 2 (3.8%) | 1.0000<sup>3</sup> | 3 (2.9%) | 18 (35.3%) | 8 (16.7%) | 0.0353<sup>2</sup> | 26 (26.3%) | <0.0001<sup>2</sup> |
| Urinary, n (%) | 21 (40.4%) | 33 (63.5%) | 0.0185<sup>2</sup> | 54 (51.9%) | 2 (3.9%) | 2 (4.2%) | 1.0000<sup>3</sup> | 4 (4.0%) | <0.0001<sup>2</sup> |
| Respiratory, n (%) | 2 (3.8%) | 4 (7.7%) | 0.6781<sup>3</sup> | 6 (5.8%) | 4 (7.8%) | 1 (2.1%) | 0.3632<sup>3</sup> | 5 (5.1%) | 0.8211<sup>2</sup> |
| Intra-abdominal, n (%) | 16 (30.8%) | 7 (13.5%) | 0.0335<sup>2</sup> | 23 (22.1%) | 3 (5.9%) | 8 (16.7%) | 0.0879<sup>2</sup> | 11 (11.1%) | 0.0358<sup>2</sup> |
| Skin, n (%) | 3 (5.8%) | 1 (1.9%) | 0.6176<sup>3</sup> | 4 (3.8%) | 0 (0.0%) | 11 (22.9%) | 0.0003<sup>2</sup> | 11 (11.1%) | 0.0479<sup>2</sup> |
| Surgical Site, n (%) | 1 (1.9%) | 0 (0.0%) | 1.0000<sup>3</sup> | 1 (1.0%) | 1 (2.0%) | 1 (2.1%) | 1.0000<sup>3</sup> | 2 (2.0%) | 0.6140<sup>3</sup> |
| Other, n (%) | 1 (1.9%) | 1 (1.9%) | 1.0000<sup>3</sup> | 2 (1.9%) | 10 (19.6%) | 7 (14.6%) | 0.5077<sup>2</sup> | 17 (17.2%) | 0.0002<sup>2</sup> |
| Unknown, n (%) | 7 (13.5%) | 5 (9.6%) | 0.5393<sup>2</sup> | 12 (11.5%) | 13 (25.5%) | 10 (20.8%) | 0.5835<sup>2</sup> | 23 (23.2%) | 0.0275<sup>2</sup> |
| **Concurrent infections** |  |  |  |  |  |  |  |  |  |
| CVC, n (%) | 1 (1.9%) | 1 (1.9%) | 1.0000<sup>3</sup> | 2 (1.9%) | 1 (2.0%) | 0 (0.0%) | 1.0000<sup>3</sup> | 1 (1.0%) | 1.0000<sup>3</sup> |
| Urinary, n (%) | 0 (0.0%) | 3 (5.8%) | 0.2427<sup>3</sup> | 3 (2.9%) | 7 (13.7%) | 7 (14.6%) | 0.9026<sup>2</sup> | 14 (14.1%) | 0.0038<sup>2</sup> |
| Respiratory, n (%) | 5 (9.6%) | 2 (3.8%) | 0.4367<sup>3</sup> | 7 (6.7%) | 9 (17.6%) | 5 (10.4%) | 0.3021<sup>2</sup> | 14 (14.1%) | 0.0831<sup>2</sup> |
| Intra-abdominal, n (%) | 6 (11.5%) | 2 (3.8%) | 0.2695<sup>3</sup> | 8 (7.7%) | 5 (9.8%) | 1 (2.1%) | 0.2056<sup>3</sup> | 6 (6.1%) | 0.6465<sup>2</sup> |
| Skin, n (%) | 1 (1.9%) | 0 (0.0%) | 1.0000<sup>3</sup> | 1 (1.0%) | 1 (2.0%) | 5 (10.4%) | 0.1050<sup>3</sup> | 6 (6.1%) | 0.0601<sup>3</sup> |
| Surgical Site, n (%) | 0 (0%) | 0 (0%) |  | 0 (0%) | 0 (0%) | 0 (0%) |  | 0 (0%) |  |
| Other, n (%) | 4 (7.7%) | 1 (1.9%) | 0.3627<sup>3</sup> | 5 (4.8%) | 6 (11.8%) | 7 (14.6%) | 0.6781<sup>2</sup> | 13 (13.1%) | 0.0370<sup>2</sup> |
| Required ICU, n (%) | 18 (34.6%) | 12 (23.1%) | 0.1941<sup>2</sup> | 30 (28.8%) | 17 (33.3%) | 9 (18.8%) | 0.0994<sup>2</sup> | 26 (26.3%) | 0.6806<sup>2</sup> |
| ID consult within 72h, n (%) | 15 (28.8%) | 11 (21.2%) | 0.3650<sup>2</sup> | 26 (25.0%) | 30 (58.8%) | 30 (62.5%) | 0.7083<sup>2</sup> | 60 (60.6%) | <0.0001<sup>2</sup> |

<sup>1</sup>Wilcoxon <sup>2</sup>Chi-Square <sup>3</sup>Fisher Exact
p value<sup>\*</sup>: test between pre-BCID and post-BCID for Gram-Negative
p value<sup>\*\*</sup>: test between pre-BCID and post-BCID for Gram-Positive
p value<sup>\*\*\*</sup>: test between Gram-Negative and Gram-Positive
Data represented as median (IQR), unless otherwise specified. **Abbreviations:** CCI, Charlson comorbidity index; CKD, chronic kidney disease; COPD, chronic obstructive pulmonary disease; CVA/TIA, cerebrovascular accident/transient ischemic attack; CVC, central venous catheter; DM, diabetes mellitus; HIV/AIDS, human immunodeficiency virus/acquired immunodeficiency syndrome; ICU, intensive care unit; ID, infectious disease; PVD, peripheral vascular disease.

### Microbiology Characteristics

Overall, 48.4% were gram-positive and 51.6% were gram-negative BSIs. The most frequently encountered organisms were Staphylococcus aureus, Staphylococcus epidermidis, Escherichia coli and Klebsiella pneumoniae. The distribution is shown in Table 2. The overall median time from collection to culture positivity was 12.0h (IQR: 10-16), median time from positivity to identification was 21.7h (IQR: 3.4-29.7) and median time from positivity to antimicrobial susceptibility testing (AST) was 49.5h (IQR: 43.5-55.3). The median time to culture positivity (p=0.092) and median time to susceptibility (p=0.061) were not significantly different between all four cohorts. As expected, there was a significant difference in the time to organism identification between pre- and post-BCID cohorts (27.1h vs. 3.3h, p<0.0001). Gram negative blood stream infections (GN-BSI) had shorter median times to identification (14.5h versus 25.5h, p<0.0001) and susceptibilities (48.3h versus 52.4h, p = 0.0085) when compared to Gram positive blood stream infections (GP-BSI). The data is summarized in Table 3.

**Table 2:** Microbiology of BSIs pre- and post-BCID

|  | **Pre-BCID**<br>**(n=107)** | **Post-BCID**<br>**(n=106)** |
| --- | --- | --- |
| Blood culture pathogens |  |  |
| Gram positive organisms (n=103) | 50 (48.5) | 53 (51.5) |
| *Bacillus cereus* | 1 | 0 |
| *Clostridium perfringens* | 1 | 1 |
| *Eggerthella lenta* | 1 | 0 |
| *Enterococcus faecalis* (not VRE) | 7 | 6 |
| *Enterococcus faecium* (not VRE) | 0 | 5 |
| *Enterococcus faecium* (VRE) | 2 | 0 |
| *Parvimonas micra* | 1 | 0 |
| *Staphylococcus aureus* (MRSA) | 6 | 6 |
| *Staphylococcus aureus* (MSSA) | 13 | 11 |
| *Staphylococcus epidermidis* | 10 | 4 |
| *Staphylococcus haemolyticus* | 1 | 0 |
| *Staphylococcus simulans* | 0 | 1 |
| *Staphylococcus spp.* (coagulase negative) | 0 | 2 |
| *Streptococcus agalactiae* | 1 | 2 |
| *Streptococcus anginosus spp.* | 1 | 2 |
| *Streptococcus equinus* | 0 | 1 |
| *Streptococcus gallolyticus* | 1 | 0 |
| *Streptococcus group G* | 0 | 1 |
| *Streptococcus mitis spp.* | 1 | 4 |
| *Streptococcus parasanguinis* | 0 | 1 |
| *Streptococcus pneumoniae* | 2 | 0 |
| *Streptococcus pyogenes* | 1 | 3 |
| *Streptococcus sanguinis* | 0 | 1 |
| *Viridans streptococcus spp.* | 0 | 2 |
| Gram negative organisms (n=110) | 57 (51.8) | 53 (48.2) |
| *Acinetobacter baumannii* | 2 | 1 |
| *Bacteroides fragilis* | 1 | 0 |
| *Campylobacter lari/subantarcticus* | 1 | 0 |
| *Citrobacter freundii complex* | 1 | 0 |
| *Citrobacter koseri* | 0 | 1 |
| *Enterobacter cloacae complex* | 2 | 5 |
| *Escherichia coli* | 22 | 20 |
| *Escherichia coli* (MDR) | 5 | 0 |
| *Klebsiella oxytoca* | 0 | 2 |
| *Roaultella ornithinolytica* | 1 | 0 |
| *Klebsiella pneumoniae* | 14 | 15 |
| *Proteus mirabilis* | 1 | 1 |
| *Pseudomonas aeruginosa* | 4 | 8 |
| *Stenotrophomonas maltophilia* | 2 | 0 |
| *Veillonella spp.* | 1 | 0 |

**Abbreviations:** MDR, multi-drug resistant; VRE, vancomycin-resistant *Enterococcus*.

**Table 3:** Antimicrobial stewardship and clinical outcomes by gram stain?

|  | Gram-Negative |  |  |  | Gram-Positive |  |  |  |  |
| --- | --- | --- | --- | --- | --- | --- | --- | --- | --- |
|  | Pre-BCID<br>(N=52) | Post-BCID<br>(N=52) | p-value* | Total<br>(N=104) | Pre-BCID<br>(N=51) | Post-BCID<br>(N=48) | p-value** | Total<br>(N=99) | p-value*** |
| Time to positivity and gram stain (T0), h | 11<br>(10-12.5) | 12<br>(10-15.5) | 0.1851<sup>1</sup> | 11.8<br>(10-14) | 13<br>(10-18) | 13<br>(11-16) | 0.8249<sup>4</sup> | 13<br>(11-17) | 0.0323<sup>1</sup> |
| Time to identification (TTID)*, h | 25.8<br>(18.9-30.3) | 2.6<br>(2.3-3.1) | <0.0001<sup>1</sup> | 14.5<br>(2.6-26.7) | 28.8<br>(23.1-45.7) | 17.4<br>(3.7-31.8) | 0.0002<sup>4</sup> | 25.5<br>(13.3-36.5) | <0.0001<sup>1</sup> |
| Time to susceptibilities (TTS)*, h | 49.2<br>(42.6-53.1) | 48.1<br>(41.4-52.1) | 0.6134<sup>1</sup> | 48.3<br>(42.1-52.5) | 52.8<br>(46.3-66.9) | 50.9<br>(44.8-79.5) | 0.7720<sup>4</sup> | 52.4<br>(44.8-74.1) | 0.0085<sup>1</sup> |
| Time to first appropriate de-escalation*, h | 42.3<br>(33.0-50.0) | 41.6<br>(34.1-50.3) | 0.6866<sup>1</sup> | 42.1<br>(33.0-50.2) | 45.6<br>(31.9-55.6) | 35.1<br>(20.2-45.8) | 0.0557<sup>4</sup> | 38.1<br>(29.2-51.3) | 0.4932<sup>1</sup> |
| Time to first appropriate escalation*, h | 11.3<br>(11.3-11.3) | 11.8<br>(11.3-36.0) | 1.0000<sup>1</sup> | 11.6<br>(11.3-36.0) | 33.8<br>(5.3-82.0) | 11.2<br>(5.2-59.5) | 0.7015<sup>4</sup> | 20.2<br>(5.3-59.5) | 0.6501<sup>1</sup> |
| Antibiotic change, T0-TTID, n (%) | 4 (7.7%) | 1 (1.9%) | 0.3627<sup>3</sup> | 5 (4.8%) | 8 (15.7%) | 10 (20.8%) | 0.5069<sup>2</sup> | 18 (18.2%) | 0.0027<sup>2</sup> |
| Antibiotic change, TTID-TTS, n (%) | 11 (21.2%) | 14 (26.9%) | 0.4912<sup>2</sup> | 25 (24.0%) | 15 (29.4%) | 17 (35.4%) | 0.5232<sup>2</sup> | 32 (32.3%) | 0.1892<sup>2</sup> |
| Antibiotic, post TTS, n (%) | 30 (57.7%) | 30 (57.7%) | 1.0000<sup>2</sup> | 60 (57.7%) | 18 (35.3%) | 17 (35.4%) | 0.9898<sup>2</sup> | 35 (35.4%) | 0.0014<sup>2</sup> |
| Any de-escalation, n (%) | 38 (73.1%) | 33 (63.5%) | 0.2921<sup>2</sup> | 71 (68.3%) | 32 (62.7%) | 32 (66.7%) | 0.6833<sup>2</sup> | 64 (64.6%) | 0.5846<sup>2</sup> |
| Any escalation, n (%) | 1 (1.9%) | 5 (9.6%) | 0.2050<sup>3</sup> | 6 (5.8%) | 7 (13.7%) | 7 (14.6%) | 0.9026<sup>2</sup> | 14 (14.1%) | 0.0454<sup>2</sup> |
| No antibiotic change, n (%) | 12 (23.1%) | 12 (23.1%) | 1.0000<sup>2</sup> | 24 (23.1%) | 14 (27.5%) | 11 (22.9%) | 0.6038<sup>2</sup> | 25 (25.3%) | 0.7173<sup>2</sup> |
| Length of stay, d | 4 (3-8) | 4 (3-6) | 0.1116<sup>1</sup> | 4 (3-7) | 8 (5-17) | 6 (4.8-12.5) | 0.1441<sup>4</sup> | 7 (5-15) | <0.0001<sup>1</sup> |
| Discharge to home, n (%) | 39 (75.0%) | 43 (82.7%) | 0.3368<sup>2</sup> | 82 (78.8%) | 34 (66.7%) | 32 (66.7%) | 1.0000<sup>2</sup> | 66 (66.7%) | 0.0510<sup>2</sup> |
| Discharge to facility, n (%) | 11 (21.2%) | 7 (13.5%) | 0.2998<sup>2</sup> | 18 (17.3%) | 15 (29.4%) | 13 (27.1%) | 0.7971<sup>2</sup> | 28 (28.3%) | 0.0619<sup>2</sup> |
| In-hospital death, n (%) | 2 (3.8%) | 3 (5.8%) | 1.0000<sup>3</sup> | 5 (4.8%) | 2 (3.9%) | 3 (6.3%) | 0.6716<sup>3</sup> | 5 (5.1%) | 1.0000<sup>3</sup> |

<sup>1</sup>Wilcoxon <sup>2</sup>Chi-Square <sup>3</sup>Fisher Exact
p-value\* : test between pre-BCID and post-BCID for Gram-Negative
p-value\*\* : test between pre-BCID and post-BCID for Gram-Positive
p-value<sup>\*\*\*</sup>: test between Gram-Negative and Gram-Positive

### Antimicrobial Stewardship Outcomes

The most frequently used antibiotics for empiric therapies include vancomycin (62.3%), piperacillin-tazobactam (41.5%), levofloxacin (24.0%), cefepime (12.3%) and ceftriaxone (9.0%). Changes in antimicrobial therapy were assessed in three distinct time periods: (1) change before ID (time-to-gram-stain to time-to-identification), (2) change after ID but before AST (time-to-identification to time-to-susceptibilities) and (3) change after AST (time-to-susceptibilities to hospital discharge). For both GN-BSIs and GP-BSIs, the median time to organism ID was significantly shorter post-BCID (2.6h versus 25.8h, p<0.0001 and 17.4h versus 28.8h, p=0.0002, respectively). However, there were no significant differences in the number of antibiotic changes before ID, after ID but before AST and after AST after BCID implementation. Moreover, there were no significant differences in the times to first appropriate escalation and de-escalation. However, when comparing GN-BSIs versus GP-BSIs as a whole, there were significant differences in changes before ID (4.8% vs. 18.2%, p=0.0027) and after AST (57.7% vs. 35.4%, p=0.0014). Specifically, there were significantly more changes, especially escalations, made after gram stain for GP-BSIs and more changes, especially de-escalations, made after AST for GN-BSIs. In GP-BSIs, two de-escalations were associated with a result of mecA negative, and the rest were associated with a GN etiology. Overall, there were no significant differences in length of stay, patient disposition on discharge or in-hospital mortality. The data is summarized in Table 3.

## DISCUSSION

In this study, we sought to evaluate the clinical utility of BCID in antimicrobial stewardship for gram-positive and gram-negative BSIs. The main findings in our study were: (1) BCID significantly reduced the time to organism identification for both gram-positive and gram-negative BSIs as expected, (2) BCID did not significantly reduce the time to first appropriate antimicrobial escalation or de-escalation for either GP-BSIs or GN-BSIs, (3) providers were more likely to escalate antimicrobial therapy in GP-BSIs after gram stain and more likely to de-escalate therapy in GN-BSIs after susceptibilities, and lastly (4) while there were no significant differences in changes in antimicrobial therapy after organism identification by BCID or gram stain status, over a quarter of providers (28.1%) still made changes after organism identification.

We present one of the first studies to evaluate the impact of gram stain results as part of an outcome analysis before and after BCID implementation. Previous studies, from retrospective observation studies to a prospective randomized controlled trial, have shown benefits of the implementation of rapid BCID, particularly in conjunction with an ASP, on antimicrobial stewardship and patient outcomes. In one retrospective study by Perez et al, investigators found that rapid BCID resulted in reduced lengths of stays, reduced cost of hospitalization and reduced 30-day mortality in patients with gram-negative BSIs. In gram-positive BSIs, rapid BCID combined with ASP implementation decreased the time to targeted therapy, decreased the duration of unnecessary antibiotics, and decreased lengths of stay and hospitalization costs (3, 13).

We present one of the first studies to analyze the effect of BCID on GN-BSI and GP-BSI separately on antimicrobial decision-making and associated outcomes. In this study, we found that there were no statistically significant differences in the time to first appropriate escalation and de-escalation among GN-BSIs and GP-BSIs after BCID implementation. However, there was a strong trend for GP-BSI for a reduced time to de-escalation, which is important in the era of increased emergence of resistant organisms. In addition, there was a trend for GP-BSI toward a reduced LOS, which could provide value not only to the patient but also to the hospital and society overall with reduced use of healthcare services. For GN-BSI, Our study demonstrated an overall lack of statistically significant impact of BCID on decision-making for antimicrobial therapy as well as lack of impact on patient outcomes.

Interestingly, this study has also provided additional quantitative insight on intuitive facets of clinical practice in the treatment of BSIs, since we evaluated each step in the identification and reporting process to determine impact on clinical decisions. For example, in response to gram stain results, providers were more likely to escalate antibiotics in gram-positive BSIs. Providers were more likely to wait to de-escalate antibiotics in gram-negative BSIs until after susceptibility results, likely due to a concern for multi-drug resistant organisms.

Based on the current evidence presented in this study, there were no significant differences in the time to first appropriate antimicrobial change. However, many providers (over one-quarter) still changed antimicrobial therapy after organism identification, indicating the identification of the organism likely plays a role in tailoring antimicrobial therapies. As expected, a greater proportion of providers were more likely to change antimicrobial therapy after susceptibility results.

Further study should be performed on the utility of BCID by gram stain morphology in larger samples. Since the time to susceptibilities appears to be highly important in antimicrobial stewardship, further research and development into accurate and rapid prediction of susceptibilities may provide the most impactful information sooner. One example of this type of technology, the Accelerate Pheno test, was recently cleared by FDA and evaluated by a large group (14).

## LIMITATIONS

There were limitations to this study. Firstly, this was a retrospective study with a modest sample size. While we attempted to mitigate this with the method of cohort selection, we cannot fully exclude the possibility of bias. Likewise, the sample size does not allow for subgroup analysis, such as the impact of BICD on the treatment of certain species organisms that may require more early tailored therapy. Second, the management of BSIs was largely left to the discretion of individual providers based on their personal clinical judgment, with the ASP group providing recommendations. Third, our tertiary referral center patient population, generally older and with multiple significant medical comorbidities, may not be representative of the general population. Lastly, we were unable to account for informal consultations (“curbside” consultations) with the hospital infectious disease consultation service.

## CONCLUSION

As expected, BCID significantly reduced the time to identification for both GP-BSIs and GN-BSIs. While BCID did not significantly reduce the time to first appropriate antimicrobial escalation and de-escalation, there was a strong trend towards a clinical impact for GP-BSI, but not for GN-BSI. It is possible that the implementation of both BCID and a highly active ASP may improve clinical outcomes in both GP-BSIs and GN-BSIs. Further research on the clinical utility of BCID for specific organisms and development of more rapid methods of susceptibility prediction is warranted.

## ACKNOWLEDGEMENTS

This research received no specific grant from any funding agency in the public, commercial, or not-for-profit sectors. None of the authors has a relevant conflict to disclose. No competing interests declared. The authors wish to thank the dedicated staff of the Mayo Clinic Arizona Microbiology laboratory.

